# Genetic engineering of *Acidithiobacillus ferridurans* with CRISPR-Cas9/dCas9 systems

**DOI:** 10.1101/2022.03.14.484339

**Authors:** Jinjin Chen, Yilan Liu, Radhakrishnan Mahadevan

**Affiliations:** Department of Chemical Engineering and Applied Chemistry, University of Toronto, Toronto, Canada; Institute of Biomedical Engineering, University of Toronto, Toronto, Canada

**Keywords:** *Acidithiobacillus ferridurans*, synthetic biology, CRISPR-Cas9, CRISPR-dCas9, genome editing

## Abstract

Genus *Acidithiobacillus* includes a group of Gram-negative Fe/S-oxidizing acidophilic chemolithotrophic bacteria that are extensively studied and used for biomining processes. Synthetic biology approaches are key means to study and improve their biomining performance. However, efficient genetic manipulations in *Acidithiobacillus* are still major bottlenecks. In this study, we report a simple and efficient pAFi system (CRISPR-dCas9) and a scarless pAF system (CRISPR-Cas9) for genetic manipulations in *A. ferridurans* JAGS. The pAFi system harboring both dCas9 and sgRNA was constructed based on pBBR1MCS-2 to knockdown *HdrA* and *TusA* genes, separately, of which the transcription levels were significantly downregulated by 48% and 93%, separately. The pAF system carrying pCas9-sgRNA-homology arms was constructed based on pJRD215 to delete *HdrB3* gene and overexpress *Rus* gene. Our results demonstrated that the pAF system is a fast and efficient genome editing method with an average rate of 15-20% per transconjugant in one recombination event, compared to 10^-3^ and then 10^-2^ in two recombination events by traditional markerless engineering strategy. Moreover, with these two systems, we successfully regulated iron and sulfur metabolisms in *A. ferridurans* JAGS: the deletion of *HdrB3* reduced 48% of sulfate production, and substitution overexpression of *Rus* promoter showed 8.82-fold of mRNA level and enhanced iron oxidation rate. With these high-efficient genetic tools for *A. ferridurans*, we will be able to study gene functions and create useful recombinants for biomining applications. Moreover, these systems could be extended to other *Acidithiobacillus* strains and promote the development of synthetic biology-assisted biomining.

**Highlights:**

- Two shuttle vectors were constructed for *Acidithiobacillus ferridurans*
- All-in-one pAFi (CRISPR-dCas9) and pAF (CRISPR-Cas9) systems were built up for gene knockdown and genome editing, separately
- The transcription levels of *HdrA* and *TusA* were reduced 48% and 93% using pAFi system and thus suppressed sulfur oxidation
- *HdrB3* deletion and *Rus* overexpression were achieved using pAF system and showed significant effects on sulfur and iron oxidation respectively
- Our pAF system facilitated genome editing in *Acidithiobacillus ferridurans* with high efficiency (15-20%) in less than 4 weeks

## 1. Introduction

Genus *Acidithiobacillus* includes a group of Gram-negative chemolithoautotrophic acidophiles that are prevalent in natural and manmade extremely acidic environments such as acid mine drainage. They obtain energies by oxidizing iron- and/or sulfur, fix carbon dioxide as carbon source, and can survive at low pH (≤3) and metal-rich environments [1]. Due to their highly adaptation to extremely acidic and metal-rich environments and their abilities to produce bioleaching chemicals H^+^ and/or Fe^3+^, members of *Acidithiobacillus* are harnessed to recover metals and rare earth elements (REEs) from ores, mine tailings, electric wastes, and spent batteries [2–4]. Furthermore, they also hold the promising for future space biomining applications [5].

Although *Acidithiobacillus* strains show significant advantages in biomining process, there are still several challenges yet to be solved, such as the long growth cycle, low biomining rate, excess sulfuric acid generated, no selectivity when release metals, and so on. Synthetic biology holds a promising role in dealing with these problems. For example, Liu *et al*. [6] and *Inaba et al*. [7] overexpressed *rusticyanin* gene on plasmids in *A. ferrooxidans* and observed enhanced iron oxidation; Gao *et al*. [8] improved the bioleaching efficiency of *A. ferrooxidans* by overexpressing the quorum sensing operon (AfeI/R) on a plasmid; Inaba *et al*. found that overexpression of glutathione synthetase (*gshB*) increased iron oxidation under conditions of oxidative stress induced by chloride [9]. However, exogenous gene expression *via* plasmids is not stable in bioleaching environments thus promotes scientists to genetically engineer *Acidithiobacillus* strains on chromosome.

The conventional methods to do genome editing in *Acidithiobacillus* strains are all based on suicide plasmids (Fig. 1A-B). The mechanism is that since suicide plasmid cannot replicate in host cells, the whole plasmid carrying an antibiotic marker will integrate into the host genome by homologous recombination under antibiotic selection. Liu *et al*. created the single recombinant *ΔrecA:Km* in *Thiobacillus ferrooxidans* ATCC33020 (reclassified as *A. ferridurans* JCM18981) by marker exchange mutagenesis using a mobilizable suicide plasmid [10, 11] (Fig. 1A). Van Zyl *et al*. constructed *ΔarsB:Km* and *ΔtetH:Km* mutants in *A. caldus* and improved the editing efficiency by enriching single recombinant cells under selective pressure and re-plating on selective plates for spontaneous double recombinants [12] (Fig. 1A). Wang *et al.* achieved the first markerless gene deletion (*ΔpfkB*) in *A. ferrooxidans*. They added an I-SceI recognition site into the suicide plasmid p1, and then introduced another plasmid p2 carrying an I-SceI endonuclease gene to induce DSB (double strand break) in the I-SceI site to facilitate markerless genome editing [13] (Fig. 1B). Although, these methods have been successfully used to engineer *Acidithiobacillus* strains and several recombinants were created [14], they are labor-intensive and time-consuming (Supplementary Fig. 1). First, suicide plasmids show much lower transfer efficiency (~10^-7^) in *Acidithiobacillus* strains compared with replicable plasmids, such as pJRD215 (10^-6^-10^-3^) and pBBR1MCS-2 (10^-7^-10^-4^). Second, these methods have complicated experimental procedures with low editing efficiencies. For example, to implement the markerless gene deletion, even a skilled engineer will need at least 8 weeks at a low efficiency (10^-3^ per transconjugant of the first recombination event and 10^-2^ per transconjugant of the second recombination event) [13]. Therefore, a fast and high-efficient method is urgently needed for genome editing in *Acidithiobacillus* strains.

**Fig. 1.**
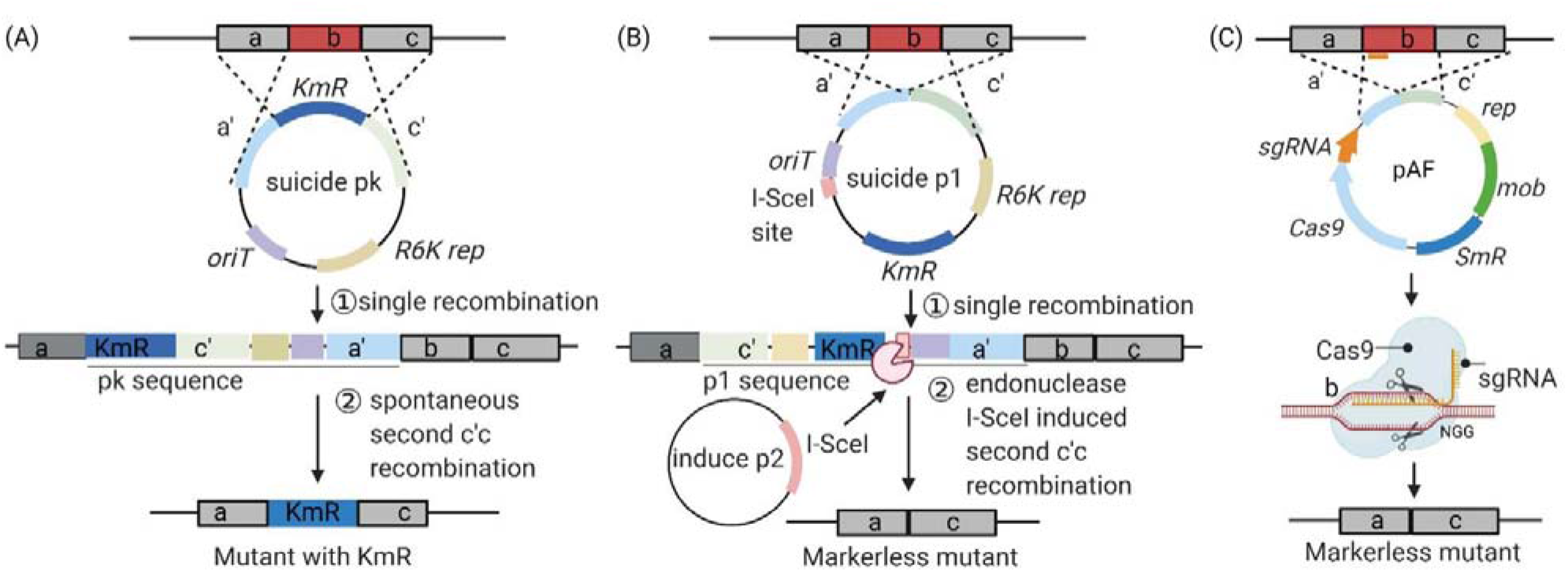
The mechanism of genome editing methods in *Acidithiobacillus* strains. The conventional methods (A-B) were based on suicide plasmids. (A) Gene deletion by marker exchange mutagenesis (Liu *et al*. 2000; van Zyl *et al*. 2008). First, the whole suicide plasmid pk inserts into the host chromosome by (1) single recombination under Km stress, and then (2) the spontaneous second recombination between c’ and c generated the mutant with b deleted. (B) Markerless gene deletion method (Wang *et al*. 2012). This method has similar mechanism as (A), except one I-SceI site (18 bp) was inserted into suicide plasmid p1 and then p2 carrying an endonuclease I-SceI gene was introduced to generate double strand breaks (DSBs) at the I-SceI site, thus induced the second recombination between c’ and c and produced the markerless mutant with b deleted. (C) Genome editing by CRISPR-Cas9 system. First, the sgRNA recognizes the target sequence b and directs Cas9 protein to recognize the PAM sequence at NGG. Second, Cas9 nuclease makes DSBs. Then the DBSs are repaired by homologous recombinant resulting in markerless mutant with b deleted.

Since 2010s, CRISPR-Cas9 system has rapidly become a powerful and robust tool for genome editing due to its easily reprogram and near universal application [15, 16]. Among these, CRISPR-Cas9, a class 2 type II CRISPR system identified in Streptococcus pyogenes composed by Cas9 and sgRNA, has rapidly become a powerful and robust tool for genome editing in different organisms due to its high orthogonality, versatility, and efficiency [17–19]. The mechanism (Fig. 1C) is that the designed sgRNA recognizes the target sequence of interested gene and directs Cas9 to recognize the NGG PAM site. Once Cas-9 finds the appropriate PAM site, its HNH and RuvC domains cleave the DNA in both strands to produce predominantly blunt-ended DSBs. Finally, the DSBs can be repaired by the host cellular DNA repair machinery, non-homologous end joining (NHEJ) or homology-directed repair (HDR) pathways. The CRISPR-Cas9 system has been applied successfully not only in model organisms like *Escherichia coli* [20] and *Saccharomyces cerevisiae* [21], but also in a variety of non-model microorganisms, such as *Lactobacillus casei* [22], *Corynebacterium glutamicum* [23], *Clostridium beijerinckii* [24] and others [25]. Furthermore, this CRISPR-Cas9 system has been successfully deployed in extremophiles [26], which encouraged us to try the CRISPR-Cas9 system in *Acidithiobacillus* strains.

In this study, we build up the pAFi (CRISPR-dCas9) and pAF (CRISPR-Cas9) systems for gene knockdown and genome editing, and successfully engineered genes involved in iron and sulfur metabolisms in *A. ferridurans*. The applicability of different plasmids and antibiotics are examined. The pAFi system-mediated gene knock down significantly reduced transcriptional levels of genes *HdrA* and *TusA* in *A. ferridurans*. We further used the pAF system for genome editing. The *HdrB3* gene was seamlessly deleted, and sulfate production was significantly reduced. The promoter of *Rus* gene was replaced with tac promoter and its expression level was enhanced 8.82-fold. Our pAFi and pAF systems provide useful tools for genetic engineering in *A. ferridurans*, thus pave the way for further synthetic biology-assisted biomining.

## 2. Materials and methods

### 2.1 Bacterial Strains, plasmids, and growth conditions

Bacterial strains and plasmids used in this study are listed in Supplementary Table 1. *E. coli* DH5α was used for plasmid construction, while *E. coli* SM10 was used as the donor to transfer plasmid into *A. ferridurans*. In this study, we used *A. ferridurans* JAGS for the genetic engineering, which we isolated from an acidic mine drainage (AMD) sample collected in September 2010 from Clarabelle Mill, near Sudbury, Ontario, Canada [27]. Its genome was sequenced and analyzed in our previous research [28]. *E. coli* was grown at 37 °C in Luria-Bertani (LB) broth or on LB agar plates. For *A. ferridurans* culture, the 9K liquid media including 9K-Fe, 9K-S and 9K-PO were used by combing 9K solution (3.0 g (NH_4_)_2_SO_4_, 0.5 g K_2_HPO_4_, 0.5 g MgSO_4_·7H_2_O and 0.1 g KCl in 1 L) [29] with either 44.22 g FeSO_4_·7H_2_O, 5 g S^0^ or 50 g pyrrhotite tailings, while 2:2 solid medium contains 2 g Na_2_S_2_O_3_·5H_2_O, 2g FeSO_4_·7H_2_O, 4.5 g (NH_4_)_2_SO_4_, 0.15 g KCl 0.75 g MgSO_4_·7H_2_O and 6 g agar in 1L. Antibiotics streptomycin (Sm) and chloramphenicol (Cm) have been used in different *Acidithiobacillus* strains with different concentrations [13, 30]. In this study, we carefully tested the Sm concentrations (100, 200 and 400 μg/mL) and Cm concentrations (10, 30 and 50 μg/mL) for recombinants screening on 2:2 solid medium. In 9K liquid medium, 400 μg/mL Sm or 50 μg/mL Cm were used to reduce plasmid loss. Absorbance of Fe^3+^ at 400 nM was recorded to monitor the continuous growth of engineered *A. ferridurans* in modified 9K-Fe medium (20 g FeSO_4_·7H_2_O in 1 L 9K solution, pH 1.5) and monitored the absorbance under 400 nm using a SPARK plate reader (TECAN) (Supplementary Fig. 2).

### 2.2 Plasmid construction

Plasmids pJRD215 and pBBR-dCas9 in our lab collections were used as the backbones to construct a series of pJRD and pBBR plasmids. The plasmid pJRD-0 and pBBR-0 were constructed for antibiotic screening and following pAFi and pAF systems construction. The functional parts for pAFi and pAF were verified as followings: the codon adaptation index (CAI) of pCas9 from *S. pyogenes* was calculated using CAI calculator (http://genomes.urv.es/CAIcal/) [31]. The promoter Ptac and Plac promoters were tested to drive pCas9 expression in *A. ferridurans*. The promoter Pj23119 was verified by driving a GFP expression and the potential sgRNA targets were designed using Cas-Designer (http://www.rgenome.net/cas-designer). A systematic BLAST search for sgRNA candidates was performed to avoid potential off-targets. DNA fragments of sgRNA and homologous arms used for plasmid construction were ordered from Twist Bioscience or constructed by one-step fusion PCR [32]. Plasmids were constructed by restriction enzyme digestion of vector DNA followed by Ligation. Restriction digests were carried out using standard protocols of NEB Restriction Endonucleases. For ligation of DNA fragments, T4 DNA ligase (Life Technologies) was used according to manufacturer’s instructions. ClonExpress MultiS One Step Cloning Kit (Vazyme, cat. #C113-01) was used for difficult constructions. Polymerase chain reaction (PCR) was performed using KAPA2G Fast/HiFi HotStart ReadyMix kit (Kapa Biosystems). Plasmids and DNA were purified using Monarch kits (NEB) according to the manufacturer’s instructions.

### 2.3 Conjugation

The constructed plasmids were transferred from *E. coli* SM10 to *A. ferridurans* JAGS using a modified conjugation method reported by Wang *et al*. [13]. In brief, the donor cells *E. coli* SM10 with plasmids were cultured at 37 °C in LB broth in presence of proper antibiotics to the late exponential growth phase. The *A. ferridurans* cells were grown in 9K-S^0^ medium to the stationary phase. The donor and recipient cells were collected by centrifugation and washed twice with basal salt solution (BSS, 4.5 g (NH_4_)_2_SO_4_, 0.15 g KCl 0.75 g MgSO_4_·7H_2_O in 1 L ddH_2_O) at 4 °C or on ice. Then donor and recipient cells were mixed in a ratio of 1:1-1:4 (total in 0.1 mL, approximately 10^9^-10^10^ cells) and spotted on a filter paper placed on a 2:2 mating plate (0.075 g FeSO_4_·7H_2_O, 2 g Na_2_S_2_O_3_·5H_2_O, 0.5 g yeast extract, 6 g agar in 1 L, pH 4.6-4.8) and incubated at 30 °C. After 2 days for plasmid transformation or 5 days for expected recombination events, the cells on the filter were resuspended and washed with BSS once, centrifuged, dissolved in 6% betaine, properly diluted, and plated on 2:2 selective plates with appropriate antibiotics at 30 °C until colonies appeared. The transfer frequencies of plasmids were calculated based on the number of transconjugants on selective plates divided by the number of recipients on nonselective plates [33].

### 2.4 Western blots

Whole cell lysis was separated on a 12% SDS-PAGE gel. Sample amount was equaled base on culture OD. Proteins on gel were transferred to a nitrocellulose membrane and detected by a Cas9 Antibody (7A9-3A3) (Novus Biotechnology, NA, USA) from mouse, and goat antimouse conjugated to HRP as a secondary antibody (Bio-Rad) followed by colorimetric detection using the Opti-4CN Kit (Bio-Rad).

### 2.5 Real-Time qPCR

A 5 mL aliquot of *A. ferridurans* culture was harvested by centrifugation at 20000 g for 5 min and then washed with BBS once. The total RNA was extracted using a general RNA extraction kit (GeneBio) following the manufacturer’s protocol. The extracted RNA was used for RT-qPCR on a CFX 96 Real-Time System (Bio-Rad) by GB-AmpTM Universal SYBR qPCR Master Mix (GeneBio). The expression levels of target genes were normalized to that of the reference gene 16S rRNA.

### 2.6 Analytical techniques

The concentration of SO_4_^2-^ in 9K-S media that cultured wild-type and engineered cells was measured using a simple turbidimetric method [34].method, of which broad-host-range The detailed experimental steps and standard curve of sulfate can be found in supplementary Fig. 3.

## 3. Results and discussion

### 3.1 Determination of Shuttle Vectors and Antibiotics for *A. ferridurans*

Several plasmids have been used as shuttle vectors from donor *E. coli* to recipient *Acidithiobacillus* cells *via* conjugation method, of which broad-host-range plasmids pJRD215 and pBBR1MCS-2 showed best performance such as transfer efficiency and stability [33, 35]. Different antibiotic selection markers including kanamycin (KmR), streptomycin (SmR) and chloramphenicol (CmR) have been applied in different *Acidithiobacillus* strains [30, 35]. In this study, the applicability of these plasmids and antibiotics are carefully examined in the newly isolate *A. ferridurans* JAGS because they are crucial for genetic engineering. First, we constructed pBBR-0 based on pBBR-dCas9 which is a derivative plasmid from pBBR1MCS-2 (Fig. 2A). We removed *dCas9* and replaced *KmR* with *CmR* because the donor *E. coli* SM10 has *KmR* in its chromosome [13]. Then we constructed pJRD-0 based on pJRD215 by removing the KmR and cohesive end site (COS) site (Fig. 2B). As shown in Fig. 2C-D, the minimal inhibitory concentration of Cm is > 10 μg/mL and Sm is > 100 μg/mL for wild-type *A. ferridurans* on 2:2 solid medium. After conjugation, we tested Cm and Sm resistances in *A. ferridurans* carrying pBBR-0 and pJRD-0, respectively. The results showed that 30 μg/mL of Cm and 200 μg/mL of Sm are appropriate concentrations for transformant selection on 2:2 solid media. The transfer efficiency of pBBR-0 is 2.13±0.54 × 10^-4^, while pJRD-0 is 1.42±0.40 × 10^-3^, which are higher than other reports [13, 30, 33]. This may because the sizes of these plasmids were reduced or *A. ferridurans* JAGS has a better feature to receive plasmids.

**Fig. 2.**
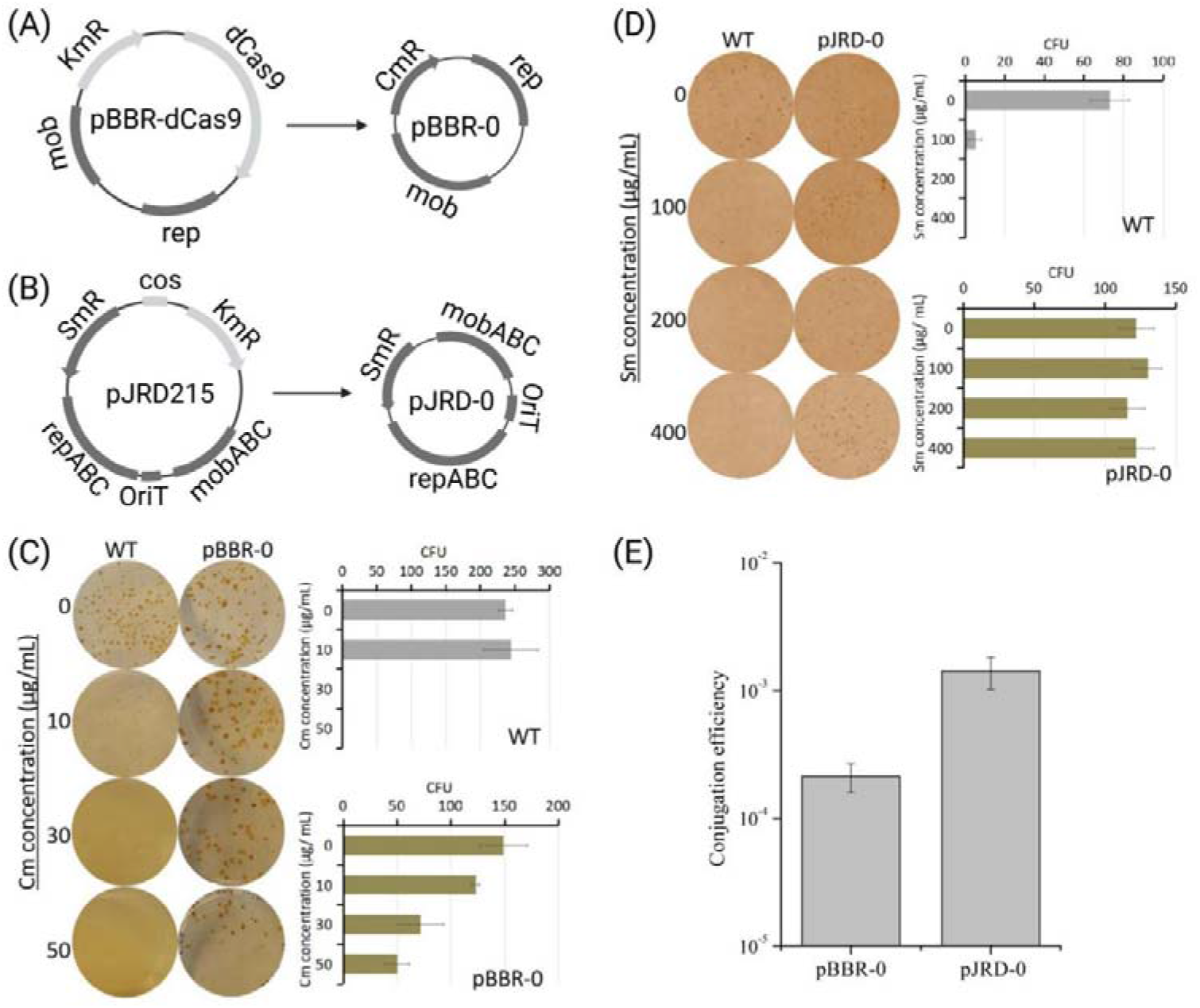
Determination of plasmids and antibiotics for *A. ferridurans* JAGS. (A-B) Construction of plasmids pBBR-0 and pJRD-0. (C-D) Chloramphenicol (Cm) and streptomycin (Sm) concentration screening for *A. ferridurans* JAGS on 2:2 solid media. (E) Transfer efficiencies of pBBR-0 and pJRD-0 from *E. coli* SM10 to *A. ferridurans* JAGS using the conjugation method.

### 3.2 Construction of pAFi and pAF systems

The plasmid pBBR-0 was used as the backbone of pAFi (CRISPR-dCas9) system and pJRD-0 with higher transfer efficiency was chosen as the backbone of pAF (CRISPR-Cas9) system (Fig. 3). First, the common bacterial promoter Pj23119 was proved to be able to drive GFP expression on pBBR-GFP in *A. ferridurans* (Fig. 3A), thus we use it to drive sgRNA transcription in *A. ferridurans*. At the same time, for pAF system construction, we first calculated the codon adaptation index (CAI) [31] of the native *Cas9* from *S. pyogenes* to predict its translation efficiency in *S. pyogenes, E. coli* and *A. ferridurans*. The CAI of native *Cas9* is 0.736 in *S. pyogenes*, 0.652 in *E. coli* and 0.591 in *A. ferridurans* which indicated that the *Cas9* might be able to express well in *A. ferridurans*. The promoter also plays an important role in gene expression. Therefore, we assigned two strong promoters Ptac and Plac, separately, to drive *Cas9* expression. Finally, we used western blot to check the Cas9 protein expression in *A. ferridurans*. As shown in Fig. 3B, the Cas9 protein from *S. pyogenes* driven by both Ptac and Plac expressed well in *A. ferridurans*. Since Ptac shows higher expression level, we used it in pAF system for the subsequent experiments. After these crucial functional parts were verified, the all-in-one pAFi system for gene knockdown and pAF system for genome editing were constructed respectively (Fig. 3).

**Fig. 3.**
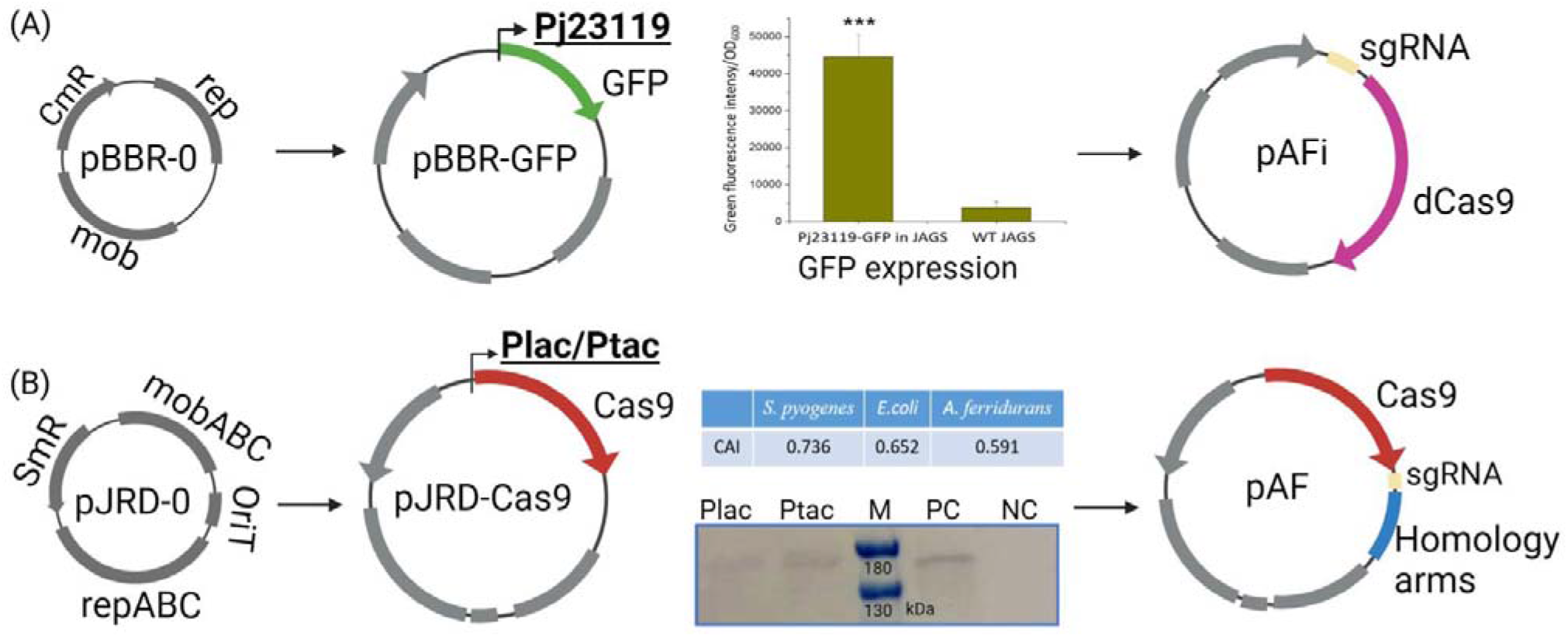
Construction of pAFi (CRISPR-dCas9) and pAF (CRISPR-Cas9) systems. (A) The pAFi was constructed based on pBBR-0. First, promoter Pj23119 was tested by driving GFP expression in *A. ferridurans* JAGS, and then the dCas9 and sgRNA parts were inserted into the plasmid formed pAFi. (B) The pAF was constructed based on pJRD-0. First, the CAI numbers of Cas9 from *S. pyogenes* were calculated in *S. pyogenes* (0.736), *E. coli* (0.652) and *A. ferridurans* (0.591) indicating good expression levels of *Cas9* in *A. ferridurans*. Second, two strong promoters Ptac and Plac were placed infront of *Cas9* gene separately. Then western-blot results demonstrated that *Cas9* from *S. pyogenes* driven by Ptac expressed well in *A. ferridurans*. Finally, the pAF was constructed by inserting *sgRNA* and homology arms of targeted gene into pJRD-PtacCas9. M, marker; PC, positive control, pCas9 expressed in *E. coli*; NC, negative control, wild-type JAGS.

### 3.3 Gene knockdown using pAFi system

Excess sulfuric acid generated by biomining process is a major source of acid mine drainage (AMD), which causes environment contamination without proper treatment [36]. Therefore, we designed pAFi system to knockdown the expression of genes involved in sulfur oxidation pathway. We chose *HdrA* (F6A13_11375) and *TusA* (F6A13_11395) in a *HdrB3-Rhd-DsrE-TusA-HdrC1B1A-orf-HdrC2B2* (F6A13_11535, 11400-11360) gene cluster as the targets because they are crucial genes responsible for sulfur trafficking, and disulfide and sulfite oxidation (Fig. 4A)[37]. We designed sgRNAs for *HdrA* and *TusA*, constructed pAFi-*dHdrA* and pAFi-*dTusA*, and conjugated them into *A. ferridurans* JAGS, separately. The wild-type *A. ferridurans* JAGS carrying pAFi-dCas9-0 (without sgRNA) was used as the control. The recombinants were first recovered in 1 mL 9K-Fe medium (Cm50). Then medium 9K-PO (Cm50) was used to culture enough cells to examine the transcriptional levels of targeted genes by qRT-PCR. As shown in Fig. 4B, the transcription levels of *HdrA* and *TusA* genes were significantly reduced by 48% and 93%, separately, compared to the control. The recovered recombinant cells were also cultured in 9K-S (Cm50) to test the effects of *HdrA* and *TusA* genes knockdown on SO_4_^2-^ production. Compared with the control, *HdrA* knockdown significantly reduced SO_4_^2-^ production but this was not observed in the recombinant of *TusA* knockdown (Fig. 4C), which is out of our expectation since TusA was reported to play a central role of cytoplasmic sulfur trafficking [38]. We hypothesized that it might be caused by plasmid loss since elemental sulfur is the only energy source in 9K-S medium, and antibiotics are not stable in low pH 9K medium. Therefore, we used PCR to test existence of pAFi-*dHdrA* and pAFi-*dTusA* in *A. ferridurans* recombinants collected from 9K-S media. The PCR results showed that pAFi-*dHdrA* was inside cells but pAFi-*dTusA* was totally lost. These results demonstrated the great capacity of the pAFi system to repress gene expression in *A. ferridurans* and the indispensable role of TusA in sulfur oxidation.

**Fig. 4.**
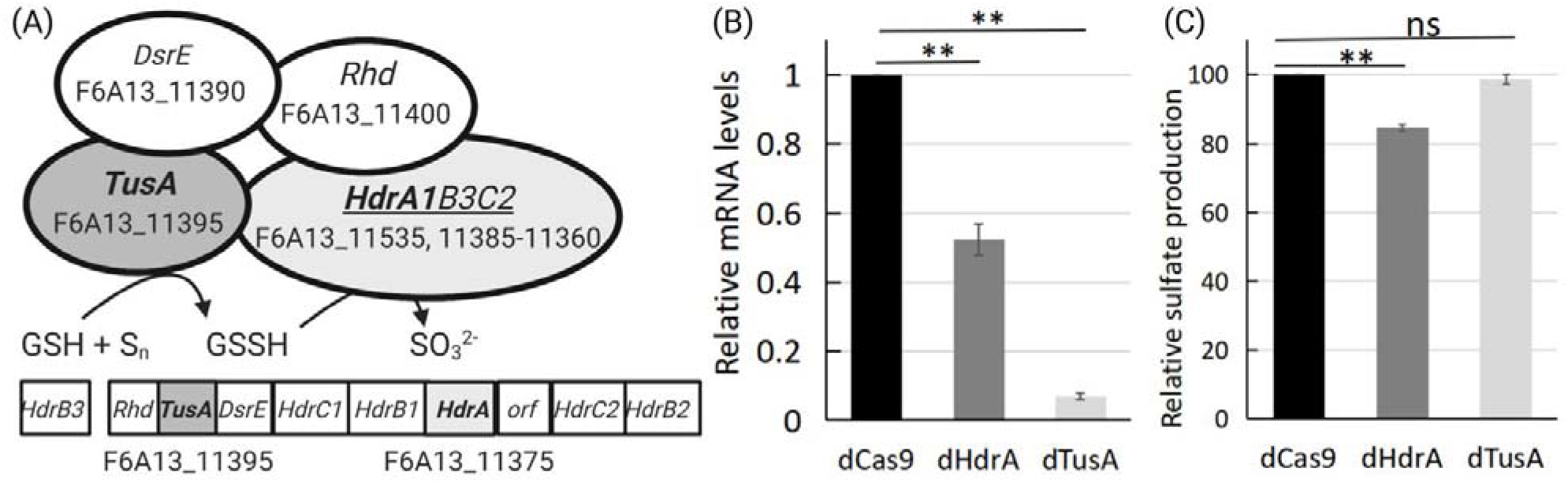
Gene knockdown in *A. ferridurans* JAGS. (A) The Hdr-like complex and Rhd-TusA-DsrE involved in sulfur oxidation in *A. ferridurans* JAGS. (B) The relative mRNA levels and (C) the relative sulfate production in dHdrA and dTus. The control strain dCas9 is wild-type *A. ferridurans* JAGS carrying pBBR-dCas9-0, while dHdrA and dTusA represent *A. ferridurans* JAGS harboring pAFi-*dHdrA* and pAFi-*dTusA*, respectively.

Recently, a similar CRISPRi system was reported in another *Acidithiobacillus* strain and they successfully knockdown a nitrogenase *nifH* gene and a Cytochrome c *cyc2* gene [39]. These indicate that CRISPRi is a powerful approach for gene repression in *Acidithiobacillus* species and would allow identification of gene functions and genetic interactions. However, plasmid stability in culture medium or natural biomining solutions is a big problem during CRISPRi application. Therefore, markerless genome editing in *Acidithiobacillus* strains is necessary for safe and stable applications in biomining industry.

### 3.4 Genome editing using the pAF system

Based on above gene knockdown results, we speculate that deletion of a gene involved in sulfur oxidation might reduce sulfuric acid generation at a certain extent. In the Hdr-like complex, the different cytoplasmic HdrB subunits harbor cysteine-rich domains which bind the unusual type [4Fe-4S] cluster and involve in disulfide reduction [37]. There are three copies of *HdrB* genes (*HdrB1*, F6A13_11380; *HdrB2*, F6A13_11360; *HdrB3*, F6A13_11535) in *A. ferridurans* JAGS (Fig. 4A). After DNA sequence alignment using PATRIC (online version 2.6.12), we found that *HdrB3* showed high sequence identity (91%) with *HdrB1*, but not *HdrB2*. We hypothesized that deletion of *HdrB3* might affect sulfur oxidation without killing the cell. Therefore, the plasmid pAF-HdrB3 (Fig. 5A) carrying a manually designed sgRNA targeting *HdrB3* (F6A13_11535) and homologous arms (~500-bp flanking *HdrB3*) was constructed and conjugated into *A. ferridurans* JAGS for deletion of *HdrB3* gene. After antibiotic screening on 2:2 solid plates, colony PCR (primers P_HdrB_1 and P_HdrB_1) and sequencing (Fig. 5A-C) were carried out and 3 recombinants were found in 20 colonies which represents 15% efficiency of this pAF system (Fig. 5A-C). We further tested the effects of different lengths of homology arms (250, 500, 750 and 1000 bp each) on *HdrB3* gene deletion efficiency by constructing plasmids pAF-HdrB3-250, pAF-HdrB3-500, pAF-HdrB3-750 and pAF-HdrB3-1000 (Fig. 5D). The results showed that when homology arms are 250 bp, no mutant was found. With the length of homology arms increased from 500 to 1000 bp, the editing efficiency increased from 15% to 20% (Fig. 5D). Next, to cure the pAF plasmid after genome editing, recombinant cells were continuously cultured in 9K-Fe medium without antibiotic stress. The pAF-HdrB3 plasmid was cured and the mutant strain ⍰HdrB3 was tested for sulfur oxidation in 9K-S medium. The results showed that *HdrB3* gene deletion reduced 48% of SO_4_^2-^ production when compared with wild-type strain (Fig. 5E). Our results showed that even there are three copies of *HdrB* genes, the deletion of *HdrB3* gene still significantly reduced the SO_4_^2-^ production when elemental sulfur as the sole energy source, which suggests its important role in elemental sulfur oxidation.

**Fig. 5.**
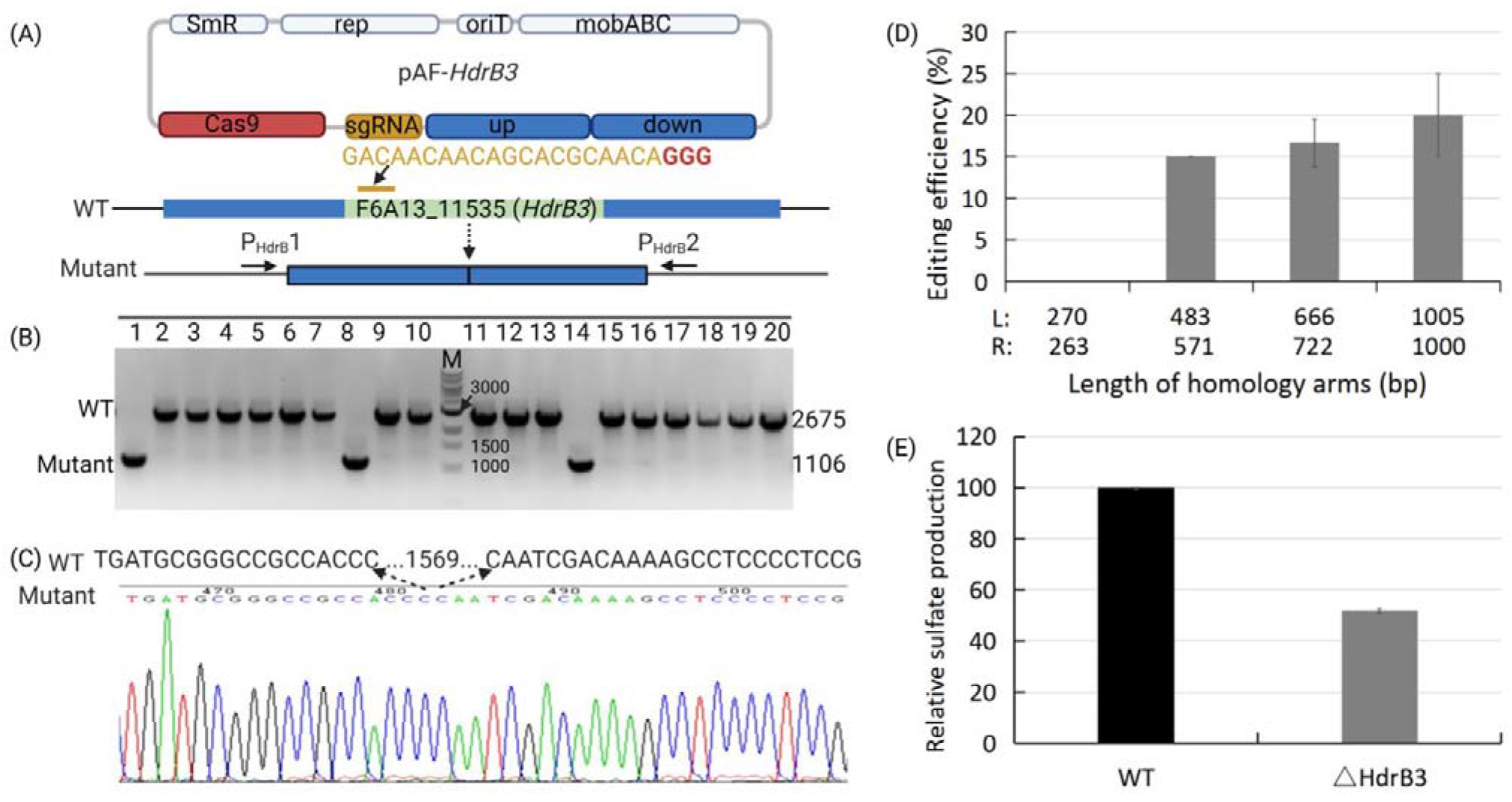
Gene deletion in *A.ferridurans* JAGS. (A-C) Design, colony PCR and sequencing result of *HdrB3* gene deletion. (D) Effects of different sizes of homologous arms on genome editing efficiency. (E) Relative sulfate production of *A.ferridurans* JAGS wild-type (WT) and HdrB3 deletion mutant (⍰HdrB3) cultured in 9K-S medium.

The strain *A. ferridurans* JAGS can obtain energy by oxidizing Fe^2+^ to Fe^3+^ and the later promotes the dissolution of pyrite. Rusticyanin *(Rus)* is a periplasmic blue copper protein, and it plays an important role in ferrous oxidation. Several attempts have been made to improve *Rus* expression in *Acidithiobacillus* strains. For examples, Liu *et al*. overexpressed *Rus* gene by introducing a wide-host-range plasmid pTRUS into *A. ferrooxidans* and they observed it not only increased the transcription levels of *Rus* gene but also other genes (*cyc1, orf, coxB, coxA*) in *Rus* operon, and resulted in improved ferrous ion oxidation rate in *A. ferrooxidans* [6]. Inaba *et al*. also overexpressed *Rus* gene using a plasmid pYI37 and noticed enhanced corrosion of stainless steel [7]. However, plasmid stability problem will hinder its industrial application. Therefore, we constructed pAF-Rus to engineer the promoter of *Rus* gene on the genome for its stable overexpression. We replaced the original promoter (Prus) with a strong constitutive promoter (Ptac) [40]. As shown in Fig. 6A, sgRNA was designed to target the upstream of Prus. The Prus and PAM site (GGG) were substituted by Ptac and non-PAM site (GTG) on mutant template. Primers P_Rus_1 and P_Rus_2 were used for colony PCR and P_Rus_3 for sequencing to verify the mutants. Fig. 6B shows the sequencing result, which demonstrates the Prus was successfully replaced by Ptac.

**Fig. 6.**
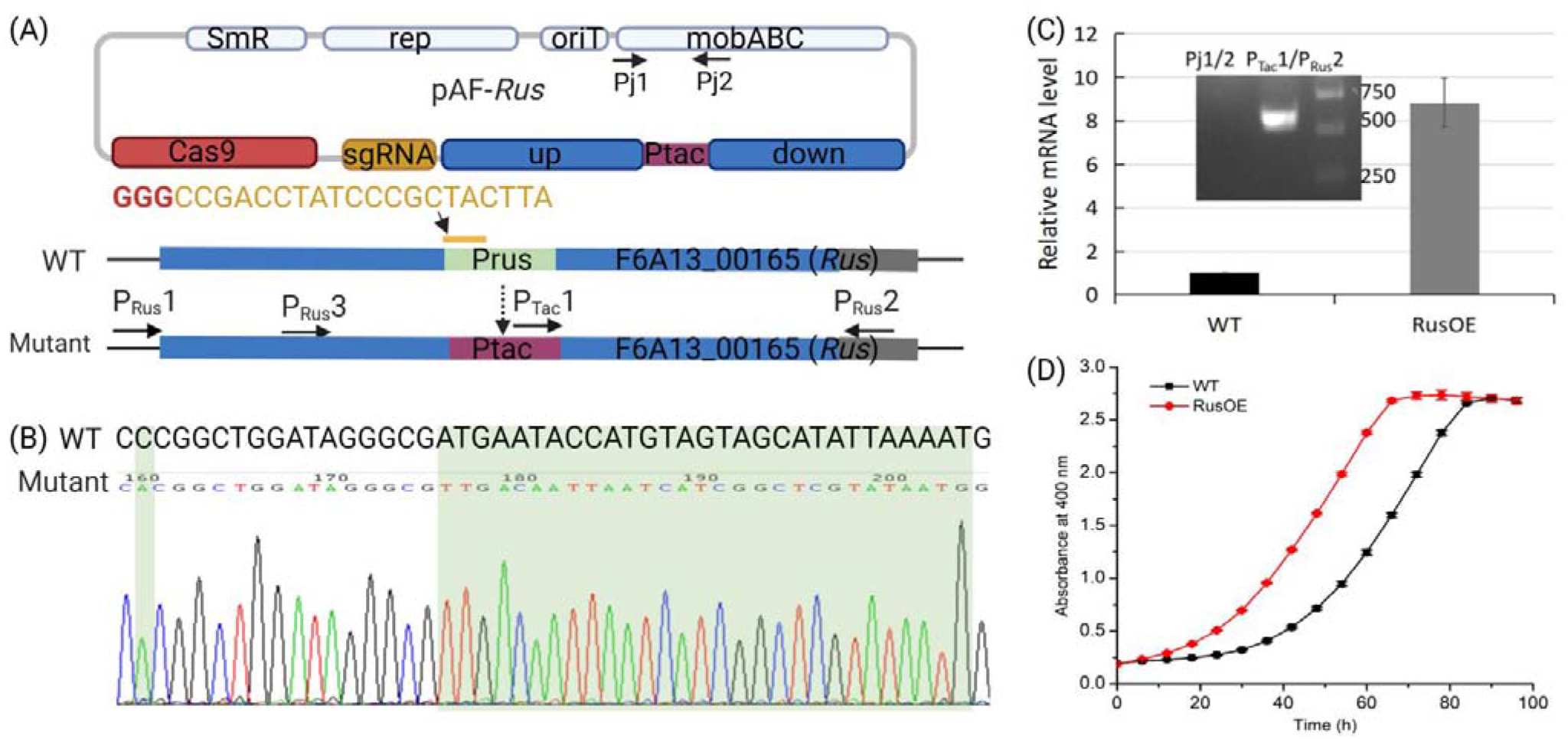
Promoter substitution assisted-gene overexpression in *A.ferridurans* JAGS. (A-B) Design and sequencing results of replacing original promoter (Prus) of *Rus* gene with Ptac. (C) The relative mRNA levels of Rus gene in *A.ferridurans* JAGS wild-type (WT) and Rus overexpression mutant (RusOE) cultured in 9K-Fe media. Gel image verified RusOE with pAF-Rus lost. Primer pairs Pj1 and Pj2 will generate a band ~ 487 bp if pAF-Rus inside the host cells, while P_Tac_1 and P_Rus_2 produce a band ~600 bp when Prus was changed to Ptac. (D) Growth curves of WT and RusOE in modified 9K-Fe medium (20 g FeSO_4_·7H_2_O in 1 L9K solution, pH 1.5) monitored by plate reader under 400 nm.

The mutant cells were recovered and continuously subcultured in 9K-Fe medium to cure the plasmid pAF-Rus. Primer pairs Pj1 and Pj2 were used to make sure pAF-Rus was lost, and primer P_Tac_1 paired with P_Rus_2 were used to confirm Ptac changed (Fig. 6A and 6C). The mutant strain without pAF-Rus inside was named as RusOE and used for following experiments. The relative mRNA level of Rus gene was tested using qRT-PCR and the result showed the transcription level of *Rus* gene was enhanced 8.2-fold in RusOE compared with wild-type strain (Fig. 6C). To monitor the growth of WT and RusOE, we cultured them in modified 9K-Fe medium (20 g FeSO_4_·7H_2_O in 1 L 9K solution, pH 1.5) in a plate reader. As shown in Fig. 6D, RusOE reached the plateau phase around 60 h after inoculation, which was much faster than the WT strain (around 80 h). Our results demonstrated that *Rus* gene can be overexpressed by changing its promoter using pAF system, and further contribute to ferrous iron oxidation.

## 4. Conclusion

In this study, we constructed two shuttle vectors for *A. ferridurans* and optimized the corresponding antibiotic concentrations for them. Then we used these two shuttle vectors to build up the CRISPRi system (pAFi) and CRISPR/Cas9 (pAF) system for genetic engineering in *A. ferridurans*. The pAFi system was successfully used to knock down *HdrA* and *TusA* genes, and qRT-PCR showed significantly reduce of their transcription levels. The pAF system was successfully used to delete *HdrB3* gene and replace promoter of *Rus* gene in the genome which led to high transcriptional level of *Rus*. Compared with traditional suicide plasmid-assisted genome editing methods, our pAF systems can achieve genome editing in a shorter period with higher efficiency in *A. ferridurans*. We envision that our pAFi/pAF systems will be broadly useful for genetic engineering in other *Acidithiobacillus* strains. Future work is needed to further optimize the efficiency for genome editing such as testing smaller Cas protein variants and more efficient sgRNA variants. Recently a minimized Cas gene, which is only 1.5 kb (cas9, around 4.1 kb) was reported for mammalian genome regulation and editing [41]. This miniature CRISPR-Cas system can be used in our pAFi/pAF plasmid systems to decrease the plasmid size, which might be able to increase conjugation and genetic editing efficiency.

## Author Contributions

JC,YL and KM conceived and designed the manuscript; JC and YL wrote the manuscript; JC, YL and KM revised the manuscript.

## Reference

1. Zhao, F. and S. Wang, Bioleaching of electronic waste using extreme acidophiles, in Electronic Waste Management and Treatment Technology. 2019, Elsevier. p. 153–174.

2. Yang, B., et al., Catalytic effect of silver on copper release from chalcopyrite mediated by Acidithiobacillus ferrooxidans. Journal of hazardous materials, 2020. 392: p. 122290.

3. Jegan Roy, J., M. Srinivasan, and B. Cao, Bioleaching as an Eco-Friendly Approach for Metal Recovery from Spent NMC-Based Lithium-Ion Batteries at a High Pulp Density. ACS Sustainable Chemistry & Engineering, 2021. 9(8): p. 3060–3069.

4. Baral, S.S., Bioleaching of rare earth elements from spent fluid catalytic cracking catalyst using Acidothiobacillus ferrooxidans. Journal of Environmental Chemical Engineering, 2021. 9(1): p. 104848.

5. Kaksonen, A.H., et al., Potential of Acidithiobacillus ferrooxidans to Grow on and Bioleach Metals from Mars and Lunar Regolith Simulants under Simulated Microgravity Conditions. Microorganisms, 2021. 9(12): p. 2416.

6. Liu, W., et al., Overexpression of rusticyanin in Acidithiobacillus ferrooxidans ATCC19859 increased Fe (II) oxidation activity. Current microbiology, 2011. 62(1): p. 320–324.

7. Inaba, Y., A.C. West, and S. Banta, Enhanced microbial corrosion of stainless steel by Acidithiobacillus ferrooxidans through the manipulation of substrate oxidation and overexpression of rus. Biotechnology and Bioengineering, 2020. 117(11): p. 3475–3485.

8. Gao, X.-Y., et al., Novel strategy for improvement of the bioleaching efficiency of Acidithiobacillus ferrooxidans based on the Afel/R quorum sensing system. Minerals, 2020. 10(3): p. 222.

9. Inaba, Y., A.C. West, and S. Banta, Glutathione synthetase overexpression in Acidithiobacillus ferrooxidans improves halotolerance of iron oxidation. Applied and Environmental Microbiology, 2021. 87(20): p. e01518–21.

10. Liu, Z., et al., Construction and characterization of arecA mutant of Thiobacillus ferrooxidans by marker exchange mutagenesis. Journal of Bacteriology, 2000. 182(8): p. 2269–2276.

11. Hedrich, S. and D.B. Johnson, Acidithiobacillus ferridurans sp. nov., an acidophilic iron-, sulfur-and hydrogen-metabolizing chemolithotrophic gammaproteobacterium. International journal of systematic and evolutionary microbiology, 2013. 63(11): p. 4018–4025.

12. van Zyl, L.J., J.M. van Munster, and D.E. Rawlings, Construction of arsB and tetH mutants of the sulfur-oxidizing bacterium Acidithiobacillus caldus by marker exchange. Applied and environmental microbiology, 2008. 74(18): p. 5686–5694.

13. Wang, H., et al., Development of a markerless gene replacement system for Acidithiobacillus ferrooxidans and construction of a pfkB mutant. Applied and environmental microbiology, 2012. 78(6): p. 1826–1835.

14. Jung, H., Y. Inaba, and S. Banta, Genetic engineering of the acidophilic chemolithoautotroph Acidithiobacillus ferrooxidans. Trends in Biotechnology, 2021.

15. Cong, L., et al., Multiplex genome engineering using CRISPR/Cas systems. Science, 2013. 339(6121): p. 819–823.

16. Doudna, J.A. and E. Charpentier, The new frontier of genome engineering with CRISPR-Cas9. Science, 2014. 346(6213).

17. Jinek, M., et al., A programmable dual-RNA-guided DNA endonuclease in adaptive bacterial immunity. science, 2012. 337(6096): p. 816–821.

18. Marraffini, L.A., The CRISPR-Cas system of Streptococcus pyogenes: function and applications. 2016.

19. Makarova, K.S., et al., Evolutionary classification of CRISPR-Cas systems: a burst of class 2 and derived variants. Nature Reviews Microbiology, 2020. 18(2): p. 67–83.

20. Zhao, D., et al., Development of a fast and easy method for Escherichia coli genome editing with CRISPR/Cas9. Microbial cell factories, 2016. 15(1): p. 1–9.

21. Stovicek, V., I. Borodina, and J. Forster, CRISPR-Cas system enables fast and simple genome editing of industrial Saccharomyces cerevisiae strains. Metabolic Engineering Communications, 2015. 2: p. 13–22.

22. Xin, Y., et al., A single-plasmid genome editing system for metabolic engineering of Lactobacillus casei. Frontiers in microbiology, 2018. 9: p. 3024.

23. Liu, J., et al., Development of a CRISPR/Cas9 genome editing toolbox for Corynebacterium glutamicum. Microbial cell factories, 2017. 16(1): p. 1–17.

24. Wang, Y., et al., Bacterial genome editing with CRISPR-Cas9: deletion, integration, single nucleotide modification, and desirable “clean” mutant selection in Clostridium beijerinckii as an example. ACS synthetic biology, 2016. 5(7): p. 721–732.

25. Selle, K. and R. Barrangou, Harnessing CRISPR-Cas systems for bacterial genome editing. Trends in microbiology, 2015. 23(4): p. 225–232.

26. !!! INVALID CITATION !!!

27. Chen, J., et al., Complete Genome Sequence of Acidithiobacillus ferridurans JAGS, Isolated from Acidic Mine Drainage. Microbiology resource announcements, 2020. 9(13).

28. Chen, J., et al., Genomic Analysis of a Newly Isolated Acidithiobacillus ferridurans JAGS Strain Reveals Its Adaptation to Acid Mine Drainage. Minerals, 2021. 11(1): p. 74.

29. Silverman, M.P. and D.G. Lundgren, Studies on the chemoautotrophic iron bacterium Ferrobacillus ferrooxidans II: Manometric Studies. Journal of bacteriology, 1959. 78(3): p. 326–331.

30. Wang, R., et al., Construction of novel pJRD215-derived plasmids using chloramphenicol acetyltransferase (cat) gene as a selection marker for Acidithiobacillus caldus. Plos one, 2017. 12(8): p. e0183307.

31. Puigbò, P., I.G. Bravo, and S. Garcia-Vallve, CAIcal: a combined set of tools to assess codon usage adaptation. Biology direct, 2008. 3(1): p. 1–8.

32. Liu, Y., J. Chen, and A. Thygesen, Efficient one-step fusion PCR based on dual-asymmetric primers and two-step annealing. Molecular biotechnology, 2018. 60(2): p. 92–99.

33. Hao, L., et al., Detection and validation of a small broad-host-range plasmid pBBR1MCS-2 for use in genetic manipulation of the extremely acidophilic Acidithiobacillus sp. Journal of microbiological methods, 2012. 90(3): p. 309–314.

34. Kolmert, Å., P. Wikström, and K.B. Hallberg, A fast and simple turbidimetric method for the determination of sulfate in sulfate-reducing bacterial cultures. Journal of microbiological methods, 2000. 41(3): p. 179–184.

35. Peng, J.-B., W.-M. Yan, and X.-Z. Bao, Plasmid and transposon transfer to Thiobacillus ferrooxidans. Journal of bacteriology, 1994. 176(10): p. 2892–2897.

36. Park, I., et al., A review of recent strategies for acid mine drainage prevention and mine tailings recycling. Chemosphere, 2019. 219: p. 588–606.

37. Wang, R., et al., Sulfur oxidation in the acidophilic autotrophic Acidithiobacillus spp. Frontiers in microbiology, 2019. 9: p. 3290.

38. Tanabe, T.S., S. Leimkühler, and C. Dahl, The functional diversity of the prokaryotic sulfur carrier protein TusA. Advances in microbial physiology, 2019. 75: p. 233–277.

39. Yamada, S., et al., Development of a CRISPR interference system for selective gene knockdown in Acidithiobacillus ferrooxidans. Journal of bioscience and bioengineering, 2021.

40. Kernan, T., A.C. West, and S. Banta, Characterization of endogenous promoters for control of recombinant gene expression in Acidithiobacillus ferrooxidans. Biotechnology and applied biochemistry, 2017. 64(6): p. 793–802.

41. Xu, X., et al., Engineered miniature CRISPR-Cas system for mammalian genome regulation and editing. Molecular Cell, 2021. 81(20): p. 4333–4345. e4.

