## supplemental files for "Genetic engineering of *Acidithiobacillus ferridurans* with CRISPR-Cas9/dCas9 systems"

**Supplementary files**

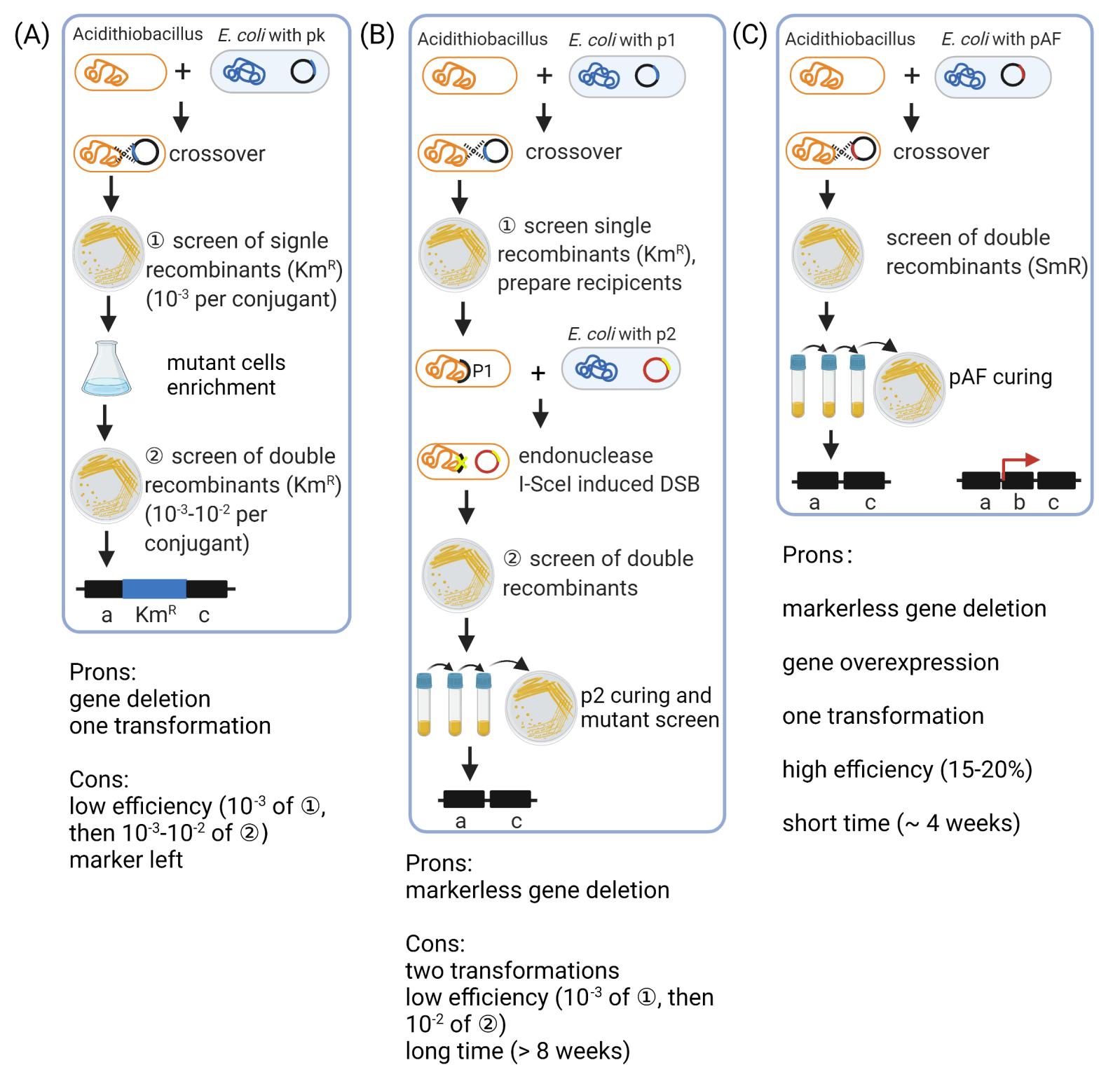

**Supplementary Fig. 1 Genome editing procedures of suicide plasmid-based genome editing methods and our CRISPR-Cas9-assisted genome editing in *Acidithiobacillus* strains.** (A) Gene deletion by marker exchange mutagenesis (Liu *et al.* 2000; van Zyl *et al.* 2008) [1, 2]; (B) Markerless gene deletion method (Wang *et al.* 2012) [3]; (C) The CRISPR-Cas9-mediated genome editing method in this study.

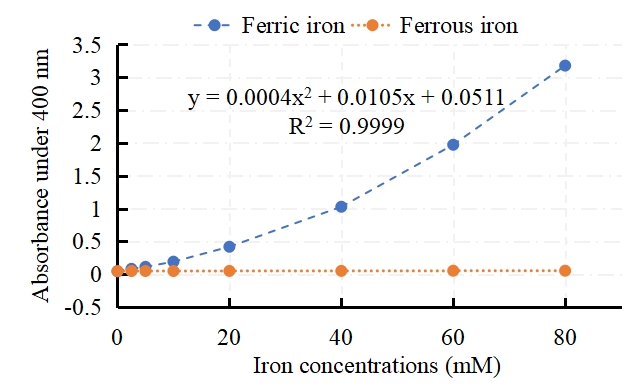

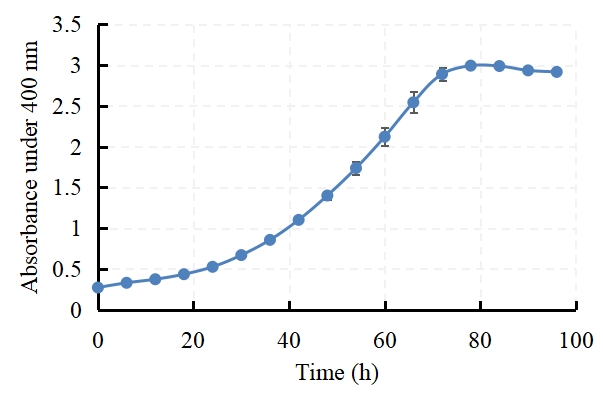

(A) **(B)**

**Supplementary Fig. 2 Method verification of using absorbance under 400 nm to monitor cell growth in adjusted 9K-Fe medium (20 g FeSO_4_·7H_2_O in 1 L 9K solution, pH 1.5)**. (A) Absorbance value under 400 nm of ferric iron and ferrous iron in 9K solution (pH 1.5) separately. Absorbance of ferrous iron can be ignored when observe the absorbance of ferric iron. (B) The growth curve of wild-type *A. ferridurans* JAGS in adjusted 9K-Fe medium. The data are given as the averages generated from three replicates.

**Supplementary**
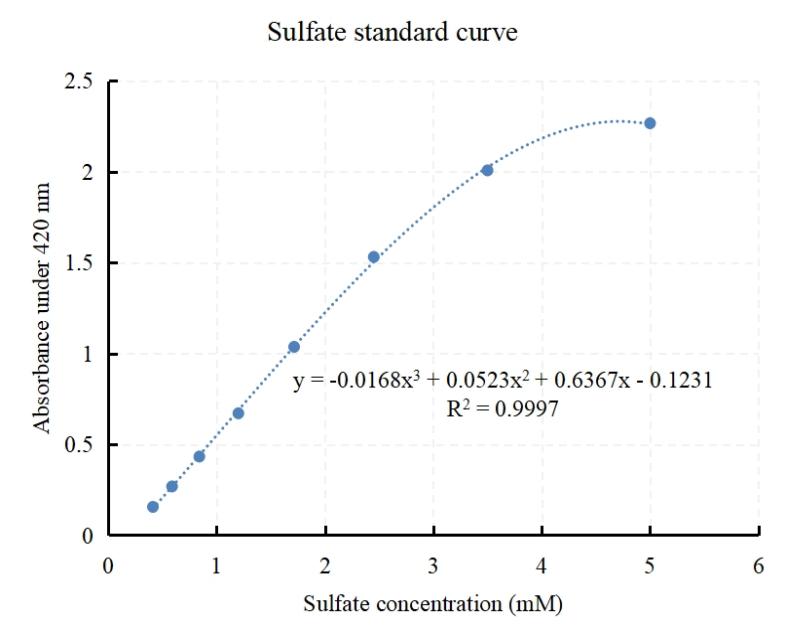
 **Fig. 3 Sulfate detection method. (Left) Standard curve and (Right) experimental steps.**

Experimental steps:

1. Reagent preparation

Reagent A: The conditioning reagent contains 150 g NaCl, 100 ml glycerol (126 g), 60 ml concentrated HCl and 200 ml 95% ethanol and is made up to 1 l with ddH_2_O.

Sample B: The sample should be diluted to fit into the range of the standard curve (0 to 5 mM).

2. Mix 1 ml of conditioning reagent A and 1 ml of sample B thoroughly in a test tube.

3. Add approximately 60 mg crushed barium chloride.

4. Mix (vortex) for 30 s.

5. Immediately pour mixture into cuvette and measure absorbance at 420 nm against a blank consisting of the complete reaction mixture excluding sulfate.

6. Compare the mean absorbance values against a standard curve made over the range of 0 to 5 mM.

7. The standard curve fitted with a third-order polynomial line.

8. Use what-if analysis- goal seek in excel to figure out the X value according to the absorbance value (Y).

Supplementary Table 1 Strains and plasmids used in this study.

| **Strains** | Description | Source |
| --- | --- | --- |
| ***E. coli*** |  |  |
| DH5α | F^-^ *endA1* *glnV44* *thi-I* *recA gyrA96 deoR nupG purB20 φ80dlacZΔM15* Δ(*lacZYA-argF*)U169, hsdR17(r_K_^–^m_K_^+^), λ^–^ | Lab collection |
| SM10 | KmR *thi-1 thr leu tonA lacY supE recA*::RP4-2–Tc::Mu, conjugation donor | [4] |
| ***A. ferridurans*** |  |  |
| WT | *A. ferridurans* JAGS wild type, isolated by our lab from an AMD in Canada | [5] |
| ΔHdrB3 | *A. ferridurans* JAGS deleted *HdrB3* gene | This study |
| RusOE | *A. ferridurans* JAGS over-expressed *rus* gene | This study |
| **Plasmids** |  |  |
| pJRD215 | SmR, KmR, IncQ, Mob^+^ | [6] |
| pJRD-0 | SmR, IncQ, Mob^+^ | This study |
| pJRD-PlacCas9 | SmR, IncQ, Mob^+^, Plac-pCas9 | This study |
| pJRD-PtacCas9 | SmR, IncQ, Mob^+^, Ptac-pCas9 | This study |
| pAF-*HdrB3* | SmR, IncQ, Mob^+^, Ptac-pCas9, Pj2319-sgRNA and HR_HdrB3_, SmR  For ***HdrB3* gene deletion** in JAGS | This study |
| pAF-*RusOE* | SmR, IncQ, Mob^+^, Ptac-pCas9, Pj2319-sgRNA and HR_Rus_, SmR  For **promoter substitution of *Rus* gene** in JAGS | This study |
| pBBR-dCas9 | KmR, pBBR1, Mob^+^, dCas9, sgRNA | Lab collection |
| pBBR-0 | CmR, pBBR1, Mob^+^ | This study |
| pBBR-dCas9-0 | CmR, pBBR1, Mob^+^, dCas9 | This study |
| pAFi-*HdrA* | CmR, pBBR1, Mob^+^, sgRNA_HdrA_, dCas9  For ***HdrA* gene knockdown** in JAGS | This study |
| pAFi-*TusA* | CmR, pBBR1, Mob^+^, sgRNA_Rhd_, dCas9  For ***TusA* gene knockdown** in JAGS | This study |

1.	Liu, Z., et al., *Construction and characterization of arecA mutant of Thiobacillus ferrooxidans by marker exchange mutagenesis.* Journal of Bacteriology, 2000. **182**(8): p. 2269-2276.

2.	van Zyl, L.J., J.M. van Munster, and D.E. Rawlings, *Construction of arsB and tetH mutants of the sulfur-oxidizing bacterium Acidithiobacillus caldus by marker exchange.* Applied and environmental microbiology, 2008. **74**(18): p. 5686-5694.

3.	Wang, H., et al., *Development of a markerless gene replacement system for Acidithiobacillus ferrooxidans and construction of a pfkB mutant.* Applied and environmental microbiology, 2012. **78**(6): p. 1826-1835.

4.	Simon, R., U. Priefer, and A. Pühler, *A broad host range mobilization system for in vivo genetic engineering: transposon mutagenesis in gram negative bacteria.* Bio/technology, 1983. **1**(9): p. 784-791.

5.	Chen, J., et al., *Complete Genome Sequence of Acidithiobacillus ferridurans JAGS, Isolated from Acidic Mine Drainage.* Microbiology resource announcements, 2020. **9**(13).

6.	Davison, J., et al., *Vectors with restriction site banks V. pJRD215, a wide-host-range cosmid vector with multiple cloning sites.* Gene, 1987. **51**(2-3): p. 275-280.
